# Effects of the Hypomethylating Agent Guadecitabine on Peripheral Blood Mononuclear Cell Methylomes and Immune Cell Populations in Small-Cell Lung Cancer Patients

**DOI:** 10.64898/2026.02.18.706553

**Authors:** Elnaz Abbasi Farid, Shu Zhang, Zhen Fu, Collin M. Coon, Daniela Matei, Shadia I. Jalal, Kenneth P. Nephew

## Abstract

**Background:** Small-cell lung cancer (SCLC) represents 15% of lung cancers and with a 5-year survival rate under 7% remains one of the deadliest malignancies. Although initially responsive to chemotherapy, rapid recurrence and resistance are common. Epigenetic modifications, particularly DNA methylation, contribute to tumor progression and therapy resistance. Guadecitabine, a hypomethylating agent (HMA), has shown promising clinical activity when combined with carboplatin in preclinical models. We evaluated the combination of guadecitabine with carboplatin as a second-line treatment for extensive-stage SCLC (NCT03913455). Here we report methylome changes in peripheral blood mononuclear cell (PBMCs) collected at baseline and during treatment from patients on the trial.

**Results:** PMBC DNA was analyzed using Infinium HumanMethylationEPIC v1.0 bead chips. Data were processed and differentially methylated positions (DMPs) were identified and analyzed for pathway enrichment using bioinformatic approaches and immune deconvolution analyses were conducted to investigate the impact on immune cell composition. Direct comparison of PBMCs between cycle 2 day 5 (C2D5; post-treatment) vs cycle 1 day 1 (C1D1; pre-treatment) revealed a greater number of hypomethylated DMPs (380 DMPs in C2D5 vs C1D1 PBMCs; p < 0.05, |β| > 20%). Moreover, when first compared with normal PBMCs from cancer-free controls, the number of hypomethylated DMPs was even greater in C2D5 than in C1D1 (1,771 vs 237 DMPs, respectively; p < 0.05, |β| > 20%). Long interspersed nucleotide elements-1 (LINE-1) were also significantly hypomethylated in PBMCs after HMA treatment (C2D5), compared to C1D1. Pathway analysis of hypomethylated DMPs revealed significant alterations in key signaling pathways including NF-κB, Rho GTPase, pulmonary fibrosis, and p75 NTR in C1D1 vs C2D5. When normal PBMCs were compared to C1D1 PBMCs, changes in IL-3 signaling, Fcγ receptor-mediated phagocytosis, and molecular mechanisms of cancer were observed. Deconvolution analysis revealed a significantly higher percentage of monocytes in C1D1 PBMCs vs normal PBMCs. However, after HMA treatment, percentages of monocytes and B cells decreased, while eosinophil percentage increased in C1D1 compared to C2D5 PBMCs.

**Conclusion:** In the first study on the global impact of HMA treatment on PBMC methylomes in SCLC patients, DNA methylation changes associated with biological pathways related to PBMC function reveal shifts in distinct immune cell populations.

**Summary:** Methylome changes in peripheral blood mononuclear cell (PBMCs) from small cell lung cancer (SCLC) patients treated with an epigenetic therapy revealed global hypomethylation and altered cancer signaling processes associated with tumor progression, immune response, therapy resistance and significant change in the proportion of immune cells. Integrating blood-based methylation biomarkers into clinical trials of epigenetic therapy and methylomic analysis of PBMCs provides direct monitoring of treatment effects in cancer patients, which may improve patient selection and enable real-time response assessment in patients receiving hypomethylating agents.

## Introduction

Small-cell lung cancer (SCLC) represents approximately 13% of all lung cancer cases and is responsible for approximately 18,000 deaths annually in the United States (1). Globally, lung cancer remains the leading cause of cancer mortality, with SCLC contributing to over 250,000 new cases and at least 200,000 deaths annually (2). With a 5-year survival rate of less than 7%, SCLC remains one of the deadliest forms of cancer. The standard treatment for extensive-stage SCLC involves systemic therapy with platinum and etoposide, recently enhanced by the addition of immune checkpoint inhibitors targeting PD-L1 (3). Despite initial responsiveness to treatment, most patients experience rapid tumor relapse, and survival outcomes remain dismal. Current therapeutic approaches have yet to provide durable responses. Platinum resistance remains a significant barrier to improving survival outcomes (4), and no effective second-line treatment options are currently available.

Epigenetic modifications, such as DNA methylation and histone modifications, play a significant role in cancer progression, heterogeneity, and resistance to treatment (5). In SCLC, DNA hypermethylation has been shown to contribute to oncogenesis (6), disease recurrence and therapy resistance (7). Hypomethylating agents (HMAs), alone or in combination with other therapeutics, have been investigated in SCLC and other solid tumors (8, 9), focusing mainly on the effects of the drug on tumor cell DNA methylation changes. However, peripheral blood mononuclear cell (PBMCs) DNA methylation changes have also been examined in lung and other cancers as potential cancer biomarkers(10-14). The effects of HMAs on DNA methylation in PBMCs from patients with SCLC have not been examined.

A phase II single-arm trial (NCT03913455) was recently conducted evaluating the combination of the HMA agent, guadecitabine, with carboplatin as a second-line treatment for extensive-stage SCLC (ES-SCLC)(15). Patients with relapsed ES-SCLC treated with guadecitabine plus carboplatin had disease control in 39.1% of cases(15). Blood samples were collected from patients on the trial and analyzed with the objective of investigating the effects of the HMA on the DNA methylome of PBMCs. Bioinformatic analysis identified both global and gene-specific changes and alterations in signaling pathways PBMCs. Deconvolution analysis revealed an unexpected finding of altered PBMC components, including monocytes and lymphocytes, suggesting that the HMA-induced changes in biological pathways which altered immune cell populations.

## Materials and Methods

### Clinical Trial

NCT03913455 clinical trial investigated a combination therapy using guadecitabine and carboplatin in adult patients diagnosed with SCLC (15). The study protocol was approved by the Institutional Review Board (IRB) to protect participant safety and maintain compliance with the Declaration of Helsinki.

### Specimens

Blood samples were collected from 24 patients as part of NCT03913455. PBMCs were isolated as follows: 16 pre-treatment samples obtained on cycle 1, day 1 (C1D1), before administration of guadecitabine and 15 post-treatment samples collected on cycle 2, day 5 (C2D5), after administration of guadecitabine on days 1–5 and carboplatin on day 5 of each 28-day cycle (for up to two cycles). Of the samples, 20 were paired (pre- and post-treatment samples from the same patients: 1005, 1008, 1009, 1010, 10105, 10106, 10118, 1020, 1021, 1023.) and 11 were unpaired (N= 6, C1D1, N= 5 C2D2). Samples from de-identified subjects without cancer served as “normal” controls (n=20; Zen-Bio, Inc., NC, USA).

DNA extraction from PBMCs was performed by using the DNeasy Blood & Tissue Kit (Qiagen Sciences Inc., Maryland, USA), according to the manufacturer’s protocol. The DNA quality and quantity were assessed using NanoDrop 2000 spectrophotometer (Thermo Fisher Scientific, Massachusetts, USA) for purity estimates, and Qubit Fluorometer (Thermo Fisher Scientific, Massachusetts, USA) to measure DNA concentration. All samples yielded high-quality nucleic acids suitable for downstream analyses. After extraction, to differentiate between methylated and unmethylated cytosines, DNA underwent bisulfite conversion using a commercially available kit (Zymo Research, California, USA) and following the manufacture’s protocol.

Genome-wide methylation profiling was conducted using the Infinium HumanMethylationEPIC v1.0 BeadChip platform (Illumina, California, USA). This platform detects methylation levels at over 850,000 CpG sites. Assay process included DNA hybridization to bead arrays, enzymatic extension, and fluorescent staining, enabling precise measurement of methylation at individual sites.

### Bioinformatic analysis

The raw idat files were processed using “openSesame” function from SeSAMe (Signal Extraction and Summarization of Array Methylation Experiments; v1.24.0) R package, which normalizes signal intensities, corrects for dye bias, and filtered low-quality probes(16, 17). SeSAMe converts idat signals to β-value, DNA methylation level which ranges from 0 (unmethylated) to 1 (fully methylated) and employs linear models to identify DMPs between groups of interest, i.e. C2D5 vs C1D1, C1D1 vs normal PBMC, and C2D5 vs normal PBMC. A β-value threshold of 0.2, along with an adjusted P-value of <0.05 (adjusted using Benjamini-Hochberg method), was applied to identify significant changes. For comparisons between C1D1 and C2D5, because the dataset includes 20 paired and 11 unpaired samples, we performed two DMP analyses: one using only the paired samples and one using all 31 samples.

To identify functional interactions of genes differentially methylated between groups by using the following approaches were pursued: 1) Ingenuity Pathway Analysis (IPA) (QIAGEN Digital Insights, California, USA), a comprehensive database for curated molecular pathways and upstream regulator analysis; 2) KEGG, which maps genes to metabolic and signaling pathways to understand cellular processes; 3) GO, which categorizes genes by biological processes, molecular functions, and cellular components; 4) TF analysis, which identifies regulatory proteins potentially driving methylation changes and 5) WP, a collaborative platform providing additional annotation and validation of pathways.

To provide additional context, CpG sites were annotated based on genomic location relative to CpG islands, shores, shelves, and transcription start sites (TSS). CpG islands were defined as regions of DNA greater than 500 bases long, with a GC content exceeding 55% and a high observed-to-expected CpG ratio. Regions extending up to 2 kilobases (kb) upstream or downstream of these islands were labeled as CpG shores, while areas 2–4 kb away were defined as CpG shelves. Sites outside these regions were classified as "open sea." In relation to gene transcripts, CpG sites were further categorized based on their proximity to key genomic features. For example, sites within 200 bases upstream of a transcription start site were classified as TSS200, while those between 200 and 1500 bases upstream were labeled as TSS1500. Additional annotations included sites in the 5′ untranslated region (5′UTR), the first exon, the gene body, and the 3′ untranslated region (3′UTR). CpG site annotation information was obtained from sesameData (17) and Noguera-Castells et al.(18).

### Deconvolution

The R package EpiDISH (19) was used to infer the fractions of a priori known cell subtypes present in our EPIC array data. In the analysis, we used “epidish” function with the Robust Partial Correlations (RPC) method (20), using centDHSbloodDMC.m as the reference data. This reference included seven immune cell types: B cells, NK cells, CD4+ T cells, CD8+ T cells, monocytes, neutrophils, and eosinophils. Next, we used the R package glmmTMB(21) to perform a beta mixed-effects regression analysis on cell type percentages between groups of samples. Additionally, we used the R package emmeans(22) to estimate means and p-values between sample groups.

### Data availability

The data generated from this study are publicly available in the Gene Expression Omnibus (GEO) repository, hosted by the National Center for Biotechnology Information (NCBI), under the accession number GSE311697.

## Results

### Guadecitabine alters the methylation landscape in circulating immune cells

Consistent with the mechanism of action of HMAs, PBMCs from patients treated with guadecitabine showed significant changes in DNA methylation after treatment. Principal Component Analysis (PCA) of methylation profiles revealed distinct clustering among normal PBMC, C1D1 (untreated) and C2D5 (HMA post-treated) (**Fig. 1A**). PC1 and PC2 explained 46.265% and 15.336% of the variance, respectively, capturing most of the variability in the dataset. Treated samples exhibited greater dispersion, indicating heterogeneous responses to treatment, while C1D1 samples clustered more tightly, reflecting consistent baseline methylation profiles. Furthermore, 380 hypomethylated CpG positions were observed on C2D5 compared to C1D1 (**Supplemental Table 1**), and differential methylation analysis further annotated 342 genes with significantly altered CpG methylation levels in C1D1 vs C2D5 overall and 78 genes in paired pre- and post-treatment samples (P<0.05; **Fig. 1B, Supplemental Table 2)**. For C1D1 vs normal PBMCs, 196 genes had significantly altered methylation changes (P<0.05; **Fig. 1C, Supplemental Table 2**), compared to 1460 genes for C2D5 vs normal PBMCs (P<0.05; **Fig. 1D, Supplemental Table 2**). In addition, CpG methylation levels across all sites revealed distinct global DNA methylation patterns in PBMC samples. Violin plots of β-values showed a typical bimodal distribution. C2D5 displayed a modest downward shift in methylation compared to baseline C1D1(**Fig. 1E**), indicating treatment-induced global hypomethylation. Taken together, these results suggest that exposure of circulating immune cells in SCLC patients treated with guadecitabine leads to an altered global methylation landscape.

**Figure 1:**
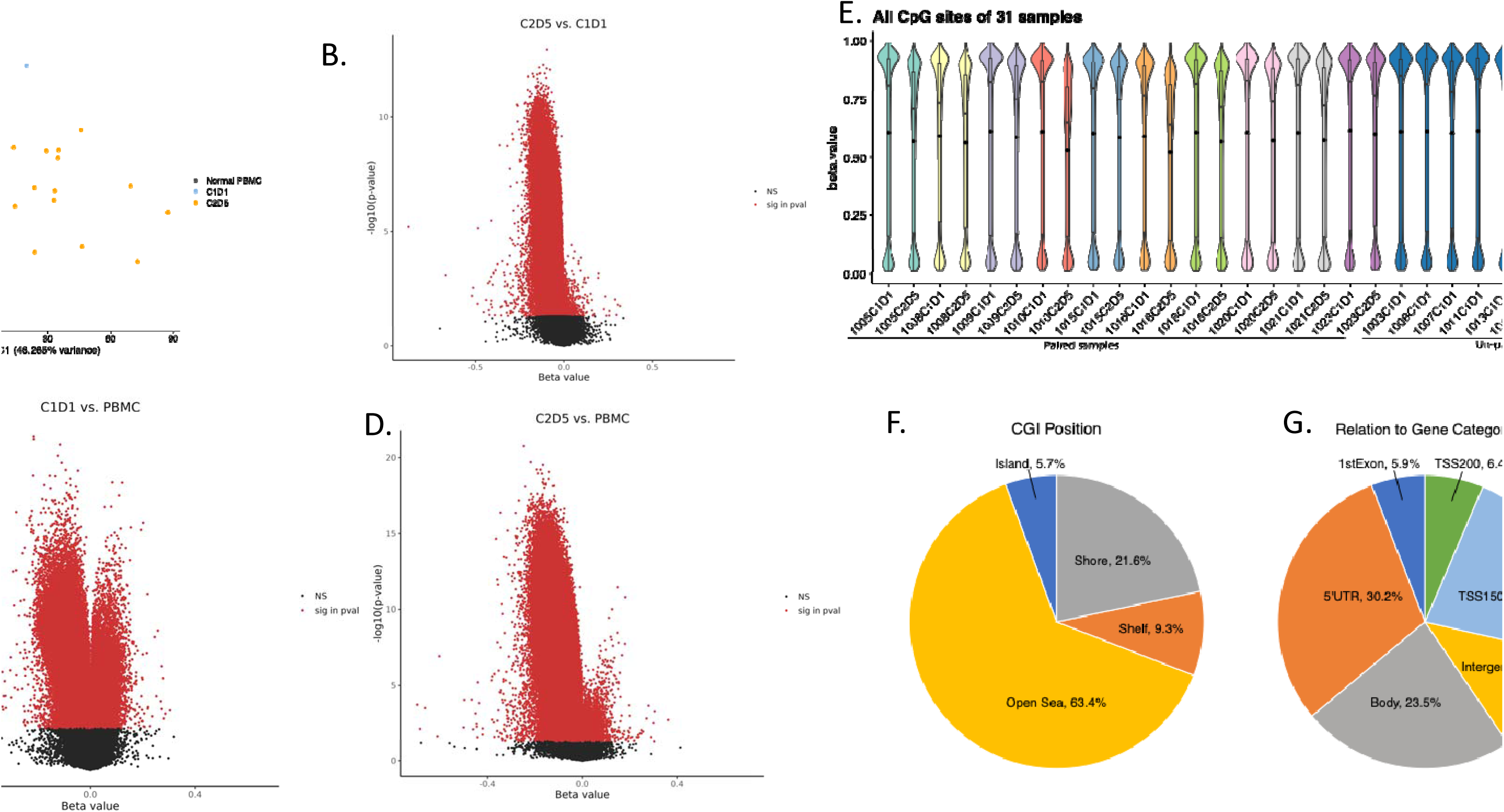
DNA methylation landscape across treated, resistant, and normal PBMCs. **(A)** Principal component analysis (PCA) of methylation data, showing separation among normal PBMC, C1D1 (untreated) and C2D1 (HMA post-treated) samples along the first two principal components. **(B)** Volcano plot comparing methylation levels between platinum-resistant and HMA-treated PBMCs. Each point represents a CpG site. Red points indicate CpGs with statistically significant differences in beta values (adjusted p < 0.05). **(C)** Volcano plot comparing platinum-resistant samples to normal PBMCs. **(D)** Volcano plot comparing HMA-treated samples to normal PBMCs. **(E)** Distribution of DNA methylation across all CpG sites in 31 PBMC samples. Each violin represents one sample, illustrating the density and spread of beta values ranging from 0 (unmethylated) to 1 (fully methylated). **(F)** Genomic annotation of DMPs relative to gene features after HMA posted treatment vs before treatment. **(G)** Pie chart showing the distribution of DMPs across CpG island contexts.

### Genomic distribution of differentially methylated positions (DMPs)

CpG probes interrogated on the Infinium HumanMethylationEPIC v1.0 BeadChip are distributed among CpG islands, which represent approximately one third of the probes; CpG island–flanking regions within 4 kb of the nearest island, referred to as shores and shelves, which account for another third of all sites; and open sea regions unrelated to CpG islands, generally located within gene bodies or intergenic regions, which comprise the remaining third of probes. DMPs were primarily found in the open sea (64%), islands (6%), shores (20%) and shelves (10%) (**Fig. 1F**). Of all guadecitabine-induced hypomethylated sites, 20% were within 1500 bp of the TSS, 6% in the first exon of a gene, while 33% was found in the 5’ UTR, and 23% of DMPs resided in gene bodies (**Fig. 1G**).

### CpG methylation levels of long interspersed nuclear element-1 (LINE-1) in PBMCs from SCLC patients

PBMCs were further assessed for global DNA methylation by analyzing LINE-1. As seen in the heatmap of β-values at LINE-1 CpG sites across all 31 PBMC samples, distinct methylation patterns were observed (**Fig. 2A**). Hierarchical clustering based on LINE-1 hypomethylation or hypermethylation showed clear segregation between before vs after HMA treatment (C1D1 vs C2D5, **Fig. 2A**). In addition, violin plots comparing CpG site methylation levels between C1D1 and C2D5 demonstrated consistent bimodal distributions, with shifts in intermediate β-value ranges (0.25–0.75) post-treatment (**Fig. 2B**). LINE-1 CpG sites showed significant methylation changes, with pre-treatment samples predominantly hypermethylated (β∼0.75) and post-treatment samples exhibiting increased hypomethylation (β ∼0.0), indicating the HMA treatment successfully induced global DNA demethylation.

**Figure 2:**
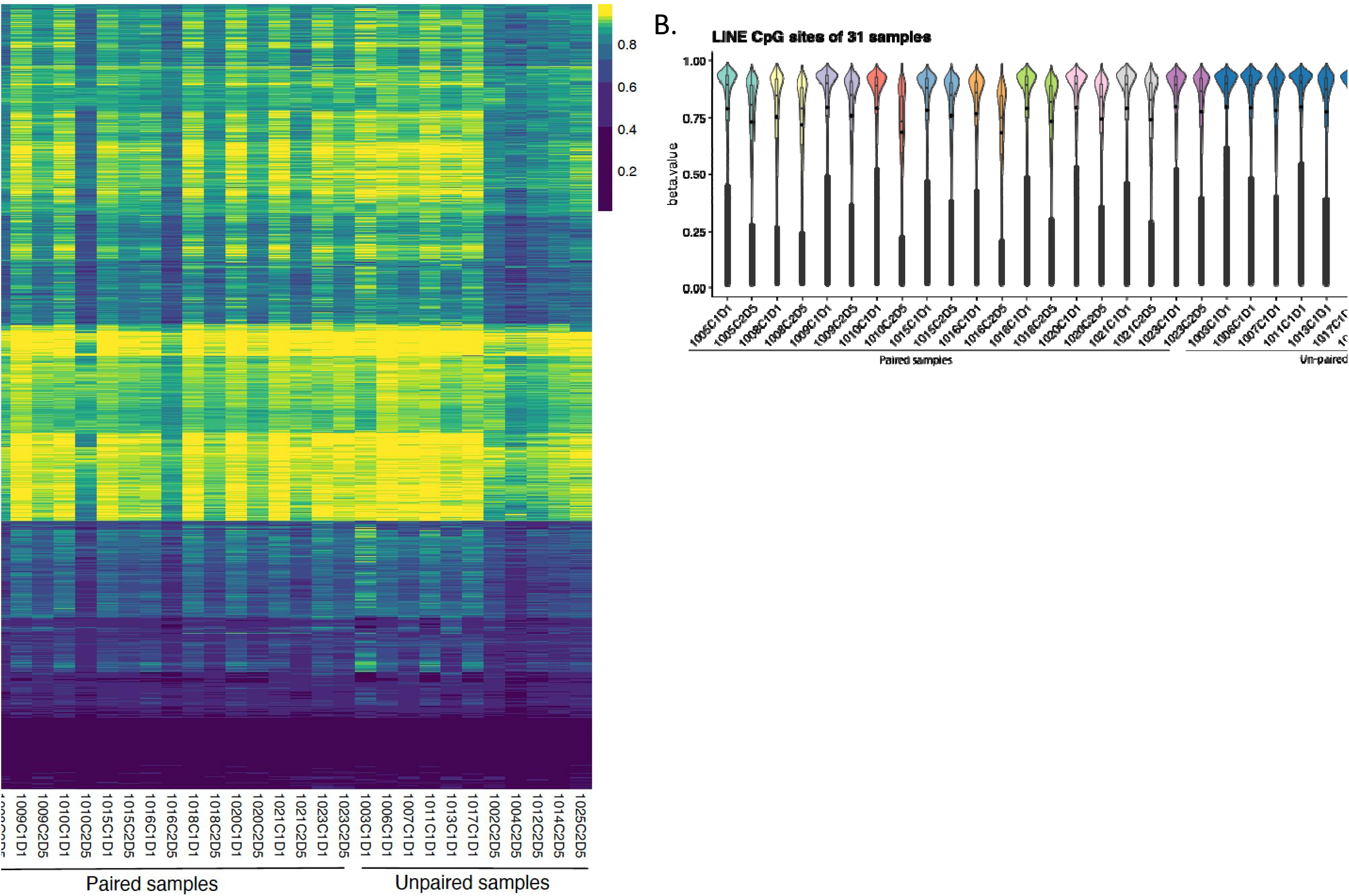
Methylation patterns of LINE-associated CpG sites in SCLC samples. **(A)** Heatmap of beta values at LINE-1 CpG sites across 31 PBMC samples. Each row represents a CpG site within LINE-1 elements, and each column represents a sample. Color intensity reflects methylation levels, with yellow indicating higher methylation and purple indicating lower methylation. Samples are hierarchically clustered based on LINE-1 methylation patterns. **(B)** Violin plot showing the distribution of beta values for LINE-1 CpG sites in each PBMC sample. Each violin represents one sample, highlighting the variation in LINE-1 methylation across the cohort. Overall trends suggest differential methylation patterns between sample groups.

### Pathway analysis of DMPs

To define pathways and identify functional interactions of genes differentially methylated between groups, exhaustive bioinformatics analyses were carried out on the DMPs. Enriched pathways in C1D1 vs normal PBMCs samples included IL-3 signaling (myeloid cell proliferation), FCγ-Receptor mediated phagocytosis (antibody-dependent cellular phagocytosis (ADCP)) and molecular mechanism of cancer (oncogenic signaling and immune evasion) (**Fig. 3A**). Enriched pathways in C2D5 vs normal PBMCs samples included RHO GTPase cycle (metastasis), opioid signaling pathway (immune modulation) and circadian rhythm pathway (regulates immune timing and cell cycle control) (**Fig. 3B**). Enriched pathways in C1D1 vs C2D5 samples included Rho GTPase signaling (metastasis), pulmonary fibrosis signaling (lung remodeling) and p75 NTR signaling (cell survival) (**Fig. 3C**). In addition, ID1 and MYC-mediated apoptosis and immunogenic cell death signaling were enriched in C2D5 v.s. C1D1 (**Supplemental Figure 1**), suggesting their relevance in therapeutic response and resistance mechanisms. The IPA network showed upregulated pathways (orange) related to immune activation and cell proliferation, driven by key regulators like STAT3 and IFNG, suggested enhanced inflammatory signaling and altered cell proliferation (**Fig. 3D**). KEGG, GO, WP, and TF databases were utilized to uncover their potential functional roles and regulatory mechanisms. As shown in **Table 1 and Supplemental Table 3**, KEGG pathway analysis highlighted the involvement of these genes in the NF-kappa B signaling, a pathway closely associated with lung cancer progression and inflammatory responses (23). GO analysis further categorized the genes into key biological processes, including developmental processes, cell differentiation, signal transduction, and tube morphogenesis, all crucial for lung tissue development and structural organization. Molecular functions such as protein binding, kinase binding, and chromatin binding were also enriched, indicating potential involvement in transcriptional regulation and intracellular signaling in SCLC.

| **Annotation Database** | **Enrichment analysis: Cancer and Immune System** |
| --- | --- |
| **KEGG** | Small cell lung cancer<br>NF-kappa B signaling |
| **GO:BP** | Regulation of Immune response,<br>Positive regulation of signal transduction |
| **GO:CC** | Cell Junction<br>Actin cytoskeleton |
| **GO:MF** | Chromatin binding<br>Kinase binding |
| **WP** | TROP2 regulatory signaling<br>Small cell lung cancer |
| **TF** | Egr-1, WT1, Sp1/Sp2/Sp6, AP-2 family, RERE, etc. |

**Figure 3:**
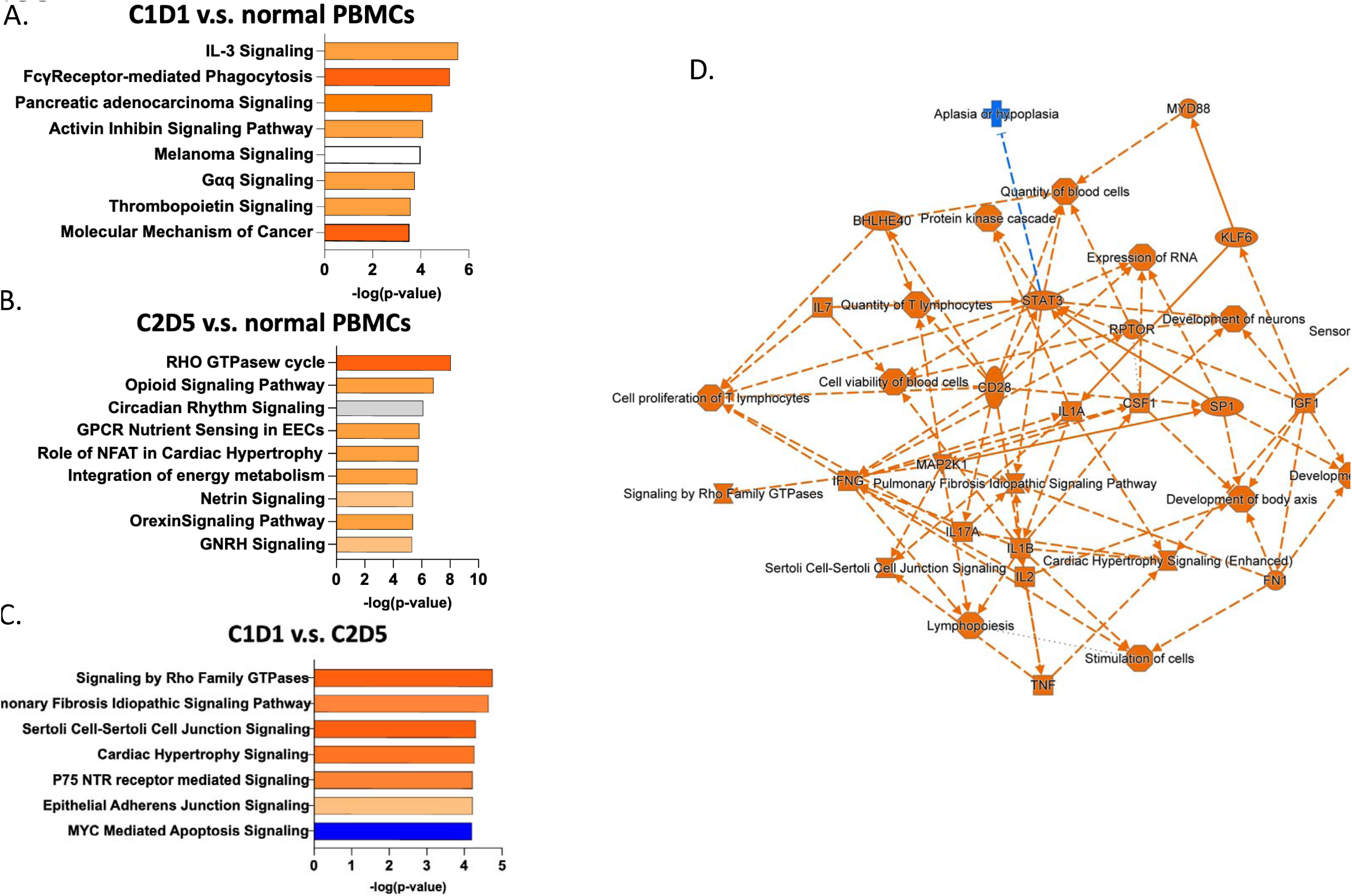
Genomic distribution and functional enrichment of differentially methylated positions (DMPs). Top canonical pathways enriched from genes annotated to DMPs **(A)** C1D1 vs normal PBMCs. **(B)** C2D5 vs normal PBMCs. **(C)** C1D1 vs C2D5 samples. **(D)** Ingenuity Pathway Analysis (IPA) network of enriched pathways derived from DMP-associated genes. Node size reflects connectivity, and orange color reflects activated pathway and blue color shows down supressed pathway.

### Immune cell composition is altered in samples from SCLC patients treated with guadecitabine

Deconvolution analysis of the methylation data demonstrated that monocytes and CD4+ were the predominant cell types in normal PBMCs and C1D1 SCLC samples (**Fig. 4A, B**). The average percentages of monocytes and neutrophils in normal vs C1D1 PBMCs were 37.52% vs 27.96%, respectively and 6.56% vs 8.02%, respectively (**Supplemental Figure 2)**. Variability in CD4+ and CD8+ T cell proportions was observed, with some samples showing reduced CD8+ T cell counts. On average, CD8+ T cell counts were 7.02% in untreated samples and 9.29% in treated samples.

**Figure 4.**
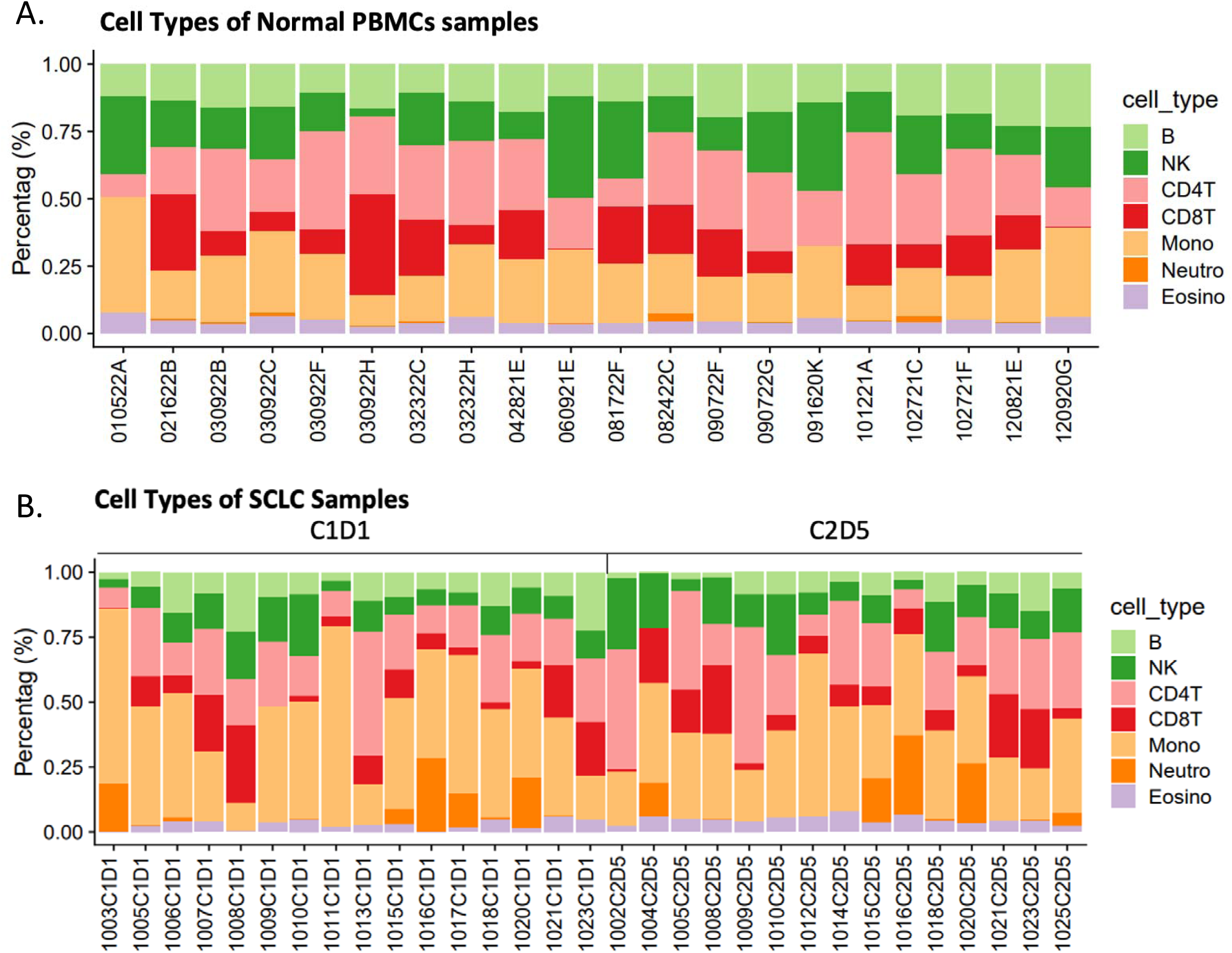
Immune cell composition in normal PBMCs and SCLC patient samples. **(A)** Stacked bar plots display the relative proportions of major immune cell types in peripheral blood mononuclear cell (PBMC) samples from non-cancerous (normal) individuals. **(B)** Immune cell proportions in PBMCs from patients with small-cell lung cancer (SCLC) before treatment (C1D1) and after treatment (C2D5), as estimated by methylation-based deconvolution. Cell types include B cells, natural killer (NK) cells, CD4+ T cells, CD8+ T cells, monocytes, neutrophils, and eosinophils, represented by distinct colors.

Several immune cell populations showed marked changes in C1D1 vs C2D5 and relative to normal PBMCs (**Table 2)**. Deconvolution analysis also revealed significant shifts in immune cell composition between comparison groups. Notably, B cells were significantly decreased between comparisons of C1D1 vs normal PBMC with odds ratios (OR) of 0.56 (p=0.001). Similarly, B-cell proportion was also lower in C2D5 compared with normal PBMCs (p < 1 × 10⁻L). Guadecitabine treatment was associated with a decreased B-cell proportion when comparing C1D1 and C2D5 samples (**Table 2**). In contrast, monocytes were significantly increased across all comparisons (C1D1 vs normal PBMCs, p = 0.0004; C2D5 vs normal PBMCs, p = 0.0037; C1D1 vs C2D5, p = 0.012), with the highest monocyte proportion observed in C1D1, suggesting an elevated myeloid compartment in PBMCs from cancer patients as well as after HMA treatment. Eosinophils significantly increased after HMA treatment (C1D1 vs C2D5, p=0.032) and were significantly lower in C1D1 compared to control PBMCs (p=0.0027). Other cell types (CD4+, CD8+ T cells, NK cells, neutrophils) exhibited directional changes but did not reach significance in all comparisons. The consistent and significant alterations in B cells and monocytes across all contrasts suggests their potential as biomarkers of disease state and treatment response in the context of HMAs.

**Table 2:** Immune Cell Type Comparisons Across PBMCs. P-values from pairwise comparisons of immune cell-type abundances across C1D1, C2D5, and PBMC samples. Significant differences (p < 0.05) suggest shifts in immune composition.

| Cell type | Comparisons | p-value |
| --- | --- | --- |
| B | C1D1 / C2D5 | **0.010725** |
| NK | C1D1 / C2D5 | 1 |
| CD4T | C1D1 / C2D5 | 0.15788 |
| CD8T | C1D1 / C2D5 | 0.847022 |
| Mono | C1D1 / C2D5 | **0.012376** |
| Neutro | C1D1 / C2D5 | 1 |
| Eosino | C1D1 / C2D5 | **0.032431** |
| B | C1D1 / PBMC | **0.001** |
| NK | C1D1 / PBMC | **0.003206** |
| CD4T | C1D1 / PBMC | 0.464751 |
| CD8T | C1D1 / PBMC | 1 |
| Mono | C1D1 / PBMC | **0.000441** |
| Neutro | C1D1 / PBMC | 0.224439 |
| Eosino | C1D1 / PBMC | **0.002684** |
| B | C2D5 / PBMC | **1E-09** |
| NK | C2D5 / PBMC | 0.51988 |
| CD4T | C2D5 / PBMC | 1 |
| CD8T | C2D5 / PBMC | 1 |
| Mono | C2D5 / PBMC | **0.003746** |
| Neutro | C2D5 / PBMC | 0.267344 |
| Eosino | C2D5 / PBMC | 1 |

## Discussion

In the current study, we demonstrate that HMA treatment induced clear differences in global hypomethylation of CpG loci in PBMCs in the setting of our previous clinical trial evaluating the combination of guadecitabine with carboplatin as a second-line treatment for SCLC (13). Pathway analysis linked hypomethylated genes in PBMCs to cancer signaling processes associated with tumor progression, immune response, and therapy resistance. In addition, we show that the proportion of monocytes, neutrophils, and T cells is highly altered by treatment with an HMA. This study is the first to investigate and demonstrate an altered methylome in PBMCs from SCLC patients treated with guadecitabine.

As DNA methylation plays a key role in SCLC progression and resistance (24), and DNA methylation profiling of PBMCs in lung cancer patients holds promise as a diagnostic tool for patient monitoring (25), PBMC methylation changes have the potential to serve as biomarkers for response to epigenetic treatments in SCLC. Furthermore, studies have associated PBMC methylation changes with tumor progression in various cancers, including breast, ovarian, colorectal, and head and neck cancers (26), suggesting potential utility in cancer monitoring and early detection. Unlike circulating free DNA (cfDNA) or circulating tumor cells (CTCs), PBMCs have a higher content in blood, are easier to extract, and offer longer DNA stability. In addition, PBMCs represent a high repeatable, minimally invasive approach with broad applicability (27), making PBMCs highly suitable for retrospective and prospective analyses.

We show significant hypomethylation of LINE-1 (Alu repetitive elements), which are heavily methylated in PBMCs (28) and thus a clinically useful pharmacodynamic marker for demethylation induced by HMAs (29). LINE-1 PBMC methylation levels clearly decreased after guadecitabine treatment, consistent with broad biological effects of guadecitabine and prior observations in trials using HMAs in patients with ovarian cancer (30, 31), and suggest the potential for impacting genomic stability and retrotransposon activity. However, whether guadecitabine-induced LINE-1 hypomethylation and presumably retrotransposon activity (32) has a direct effect on the observed changes in monocytes, neutrophils T cell proportions is unknown and will require further investigation.

In the clinical trial associated with this study (15), patients receiving second-line treatment (C2D5) had prior exposure to carboplatin (C1D1), which could contribute to methylation pattern changes. Chemotherapy-induced DNA damage has been shown to alter DNA methylation in blood, as demonstrated by Flanagan et al. (33), who also observed that similar methylation changes in both blood and tumor samples at patients’ relapse (33). Taken together, we suggest that systemic epigenetic modifications could serve as biomarkers for prognosis and treatment response either alone or in combination with tumor and/or cfDNA DNA methylation patterns.

Analysis of hypomethylated genes in PBMCs revealed cancer signaling pathways that could be relevant to the observed changes in immune cell populations. For example, Rho GTPases regulate proliferation, apoptosis, metabolism, senescence, and stemness (34). Furthermore, these processes play crucial roles in cell migration, metastasis, and interactions with the tumor microenvironment, influencing inflammation and cancer progression (35). Another specific example is an altered the pulmonary fibrosis signaling pathway. This pathway involves fibroblast and myofibroblast activation, leading to excessive ECM deposition and fibrotic foci formation, primarily driven by TGF-β, IL-17A, PDGF, and Wnt signaling (36). While typically linked to lung tissue, similar fibrotic signaling induced by guadecitabine may reflect the altered PBMC profile observed in the SCLC patients.

Based on GO and KEGG annotations, enrichment in processes related to immune signaling modulation and chromatin dynamics were observed, supporting the notion that guadecitabine-and carboplatin-induced epigenetic changes affect multiple cellular mechanisms. The findings further suggest that hypomethylation of genes involved in immune response pathways, such as ID1 and MYC-mediated apoptosis (37), may enhance tumor immunogenicity and immunogenic clearance mechanisms, providing a rationale for combining hypomethylating agents with immune checkpoint inhibitors (38). In this context, the observed increase in monocytes (6.57% to 8.02%) and reduction in neutrophils (37.52% to 27.96%) would be consistent with a shift from acute inflammation to a more regulated immune state. On the other hand, the observed reduction in CD8+ T-cell proportions in two treated samples could reflect a more immunosuppressive microenvironment, potentially counteracting the benefits of epigenetic reprogramming in some patients. Nonetheless, the findings suggest that HMA treatment induces both apoptotic and immune epigenetic remodeling responses, with potential implications for tumor cell clearance and reshaping of the tumor microenvironment. Future studies should investigate the interaction between methylation changes and immune cell dynamics to better understand potential factors influencing treatment outcomes.

We recognize that this study has limitations. The absence of tumor biopsies from these patients prevents direct correlation between PBMC and tumor methylation patterns. Epigenetic changes are known to be tissue specific, and whether DNA methylation changes in circulating PBMCs reflect tumor exposure after HMA treatment and changes in the tumor microenvironment or tumor-specific alterations in SCLC remains unknown. The lack of RNA data from PBMCs limits functional validation of how the observed genomic distribution of methylation changes (**Fig. 1**) impact gene expression in specific immune cells. As the PBMCs were not sorted, we are not able to determine mechanistically why some immune cell types responded to HMA, while others did not. It is also possible that the hypomethylation effect could be masked by an increased cytotoxic activity of carboplatin on demethylated cells.

In conclusion, these results highlight the promise of integrating blood-based methylation biomarkers into clinical trials of epigenetic therapy. Together with emerging technologies that measure epigenetic changes in cfDNA (39-41), methylomic analysis of PBMCs provides direct monitoring of treatment effects in cancer patients. Such approaches may improve patient selection and enable real-time response assessment in patients receiving HMAs. The observed epigenetic immune remodeling provides insights into HMA-induced systemic effects, including myelosuppression and immune cell modulation, warranting further investigation.

### Clinical practice points

- Epigenetic therapy with the hypomethylating agent guadecitabine combined with carboplatin as a second-line treatment in small cell lung cancer induced significant DNA hypomethylation in peripheral blood mononuclear cells, demonstrating systemic epigenetic activity.
- Pathway analysis linked hypomethylated genes in PBMCs to cancer signaling processes associated with tumor progression, immune response, and therapy resistance. In addition, the proportion of monocytes, neutrophils, and T cells was altered by treatment with an HMA, suggesting modulation of immune cell composition.
- Integrating blood-based methylation biomarkers into clinical trials of epigenetic therapy, such as methylomic analysis of PBMCs, provides direct monitoring of treatment effects in SCLC patients, may improve patient selection and enable real-time response assessment in patients receiving HMAs. The observed epigenetic immune remodeling provides insights into HMA-induced systemic effects, including myelosuppression and immune cell modulation, warranting further investigation.

## Acknowledgments

We thank Marie Adams (Genomics Core, Van Andel Institute, Grand Rapids, MI). Research funding provided by the Van Andel Research Institute - Stand Up To Cancer Epigenetics Dream Team (K.P.N.). The indicated Stand Up To Cancer Grant is administered by AACR, the scientific partner of Stand Up To Cancer. PBMC from patients with SCLC were banked and retrieved from the Pathology Core supported by National Institutes of Health, National Cancer Institute (P30CA082709-25) awarded to the IU Simon Comprehensive Cancer Center. The clinical trial was supported by Merck and Astex, Inc. K.P.N. holds the Jerry W. and Peggy S. Throgmartin Chair in Oncology is supported by P30CA082709-25.

