## Supplemental Figs 1, 2, 3 for "Effects of the Hypomethylating Agent Guadecitabine on Peripheral Blood Mononuclear Cell Methylomes and Immune Cell Populations in Small-Cell Lung Cancer Patients": SCLC_Paper_Suppl Figure1.pptx

### Slide 1
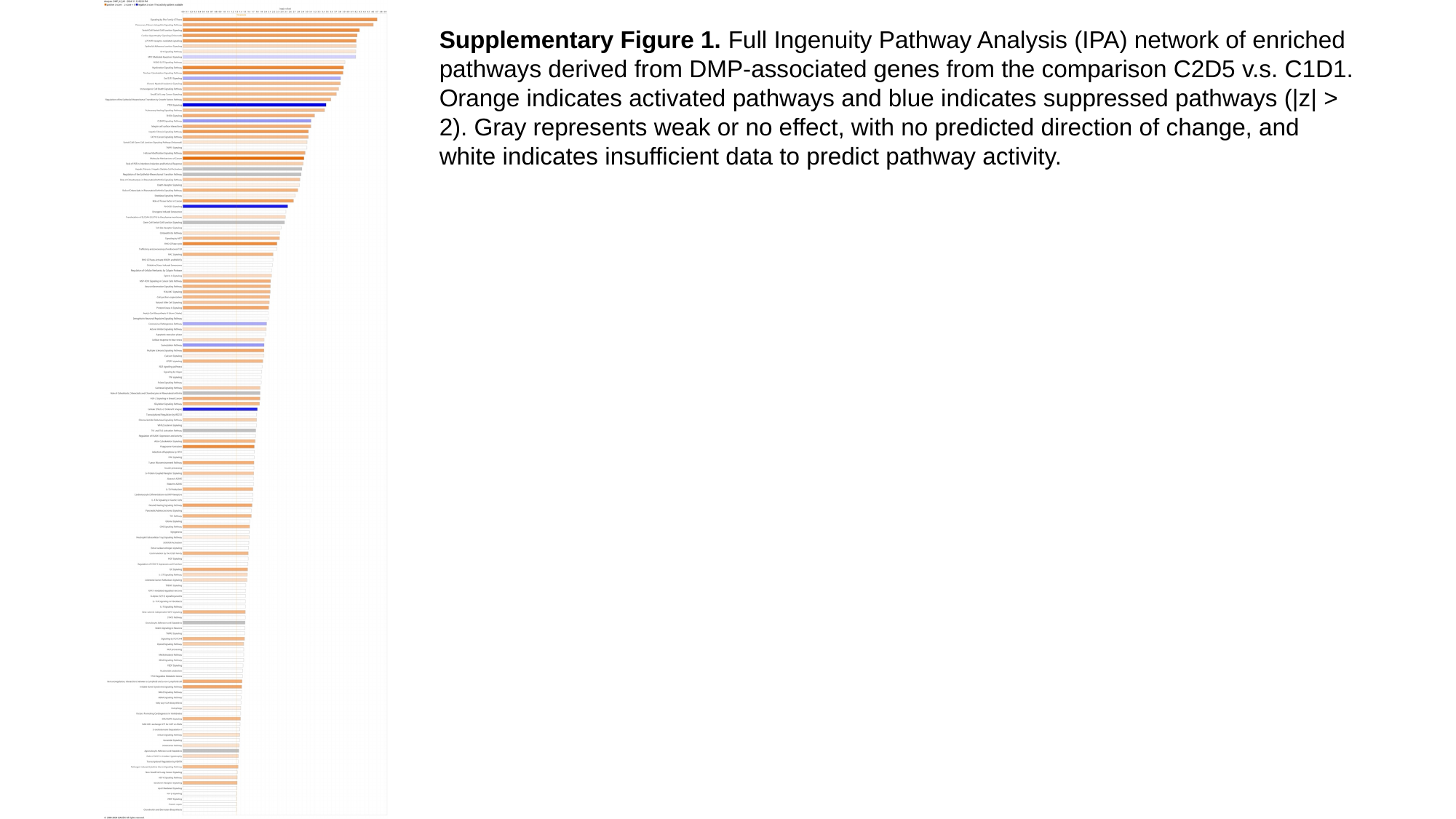

Supplementary Figure 1. Full Ingenuity Pathway Analysis (IPA) network of enriched pathways derived from DMP-associated genes from the comparison C2D5 v.s. C1D1. Orange indicates activated pathways and blue indicates suppressed pathways (|z| > 2). Gray represents weak or no effect, with no predicted direction of change, and white indicates insufficient data to predict pathway activity.
