## Supplemental Figs 1, 2, 3 for "Effects of the Hypomethylating Agent Guadecitabine on Peripheral Blood Mononuclear Cell Methylomes and Immune Cell Populations in Small-Cell Lung Cancer Patients": SCLC_Paper_Suppl Figure2.pptx

### Slide 1
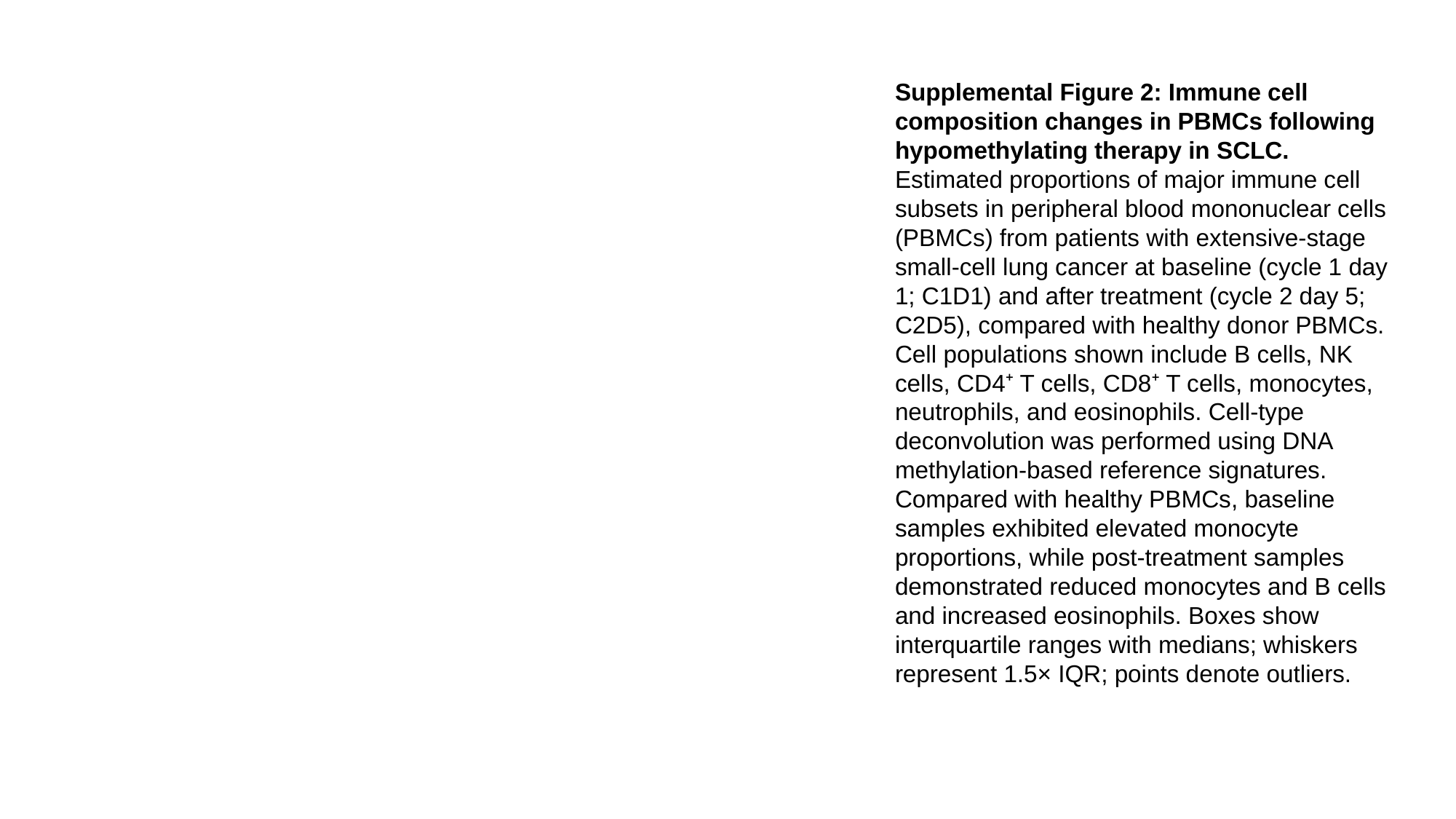

Supplemental Figure 2: Immune cell composition changes in PBMCs following hypomethylating therapy in SCLC.
Estimated proportions of major immune cell subsets in peripheral blood mononuclear cells (PBMCs) from patients with extensive-stage small-cell lung cancer at baseline (cycle 1 day 1; C1D1) and after treatment (cycle 2 day 5; C2D5), compared with healthy donor PBMCs. Cell populations shown include B cells, NK cells, CD4⁺ T cells, CD8⁺ T cells, monocytes, neutrophils, and eosinophils. Cell-type deconvolution was performed using DNA methylation-based reference signatures. Compared with healthy PBMCs, baseline samples exhibited elevated monocyte proportions, while post-treatment samples demonstrated reduced monocytes and B cells and increased eosinophils. Boxes show interquartile ranges with medians; whiskers represent 1.5× IQR; points denote outliers.
