## Supplemental Figs 1, 2, 3 for "Effects of the Hypomethylating Agent Guadecitabine on Peripheral Blood Mononuclear Cell Methylomes and Immune Cell Populations in Small-Cell Lung Cancer Patients": SCLC_Paper_Suppl Figure3_.pptx

### Slide 1
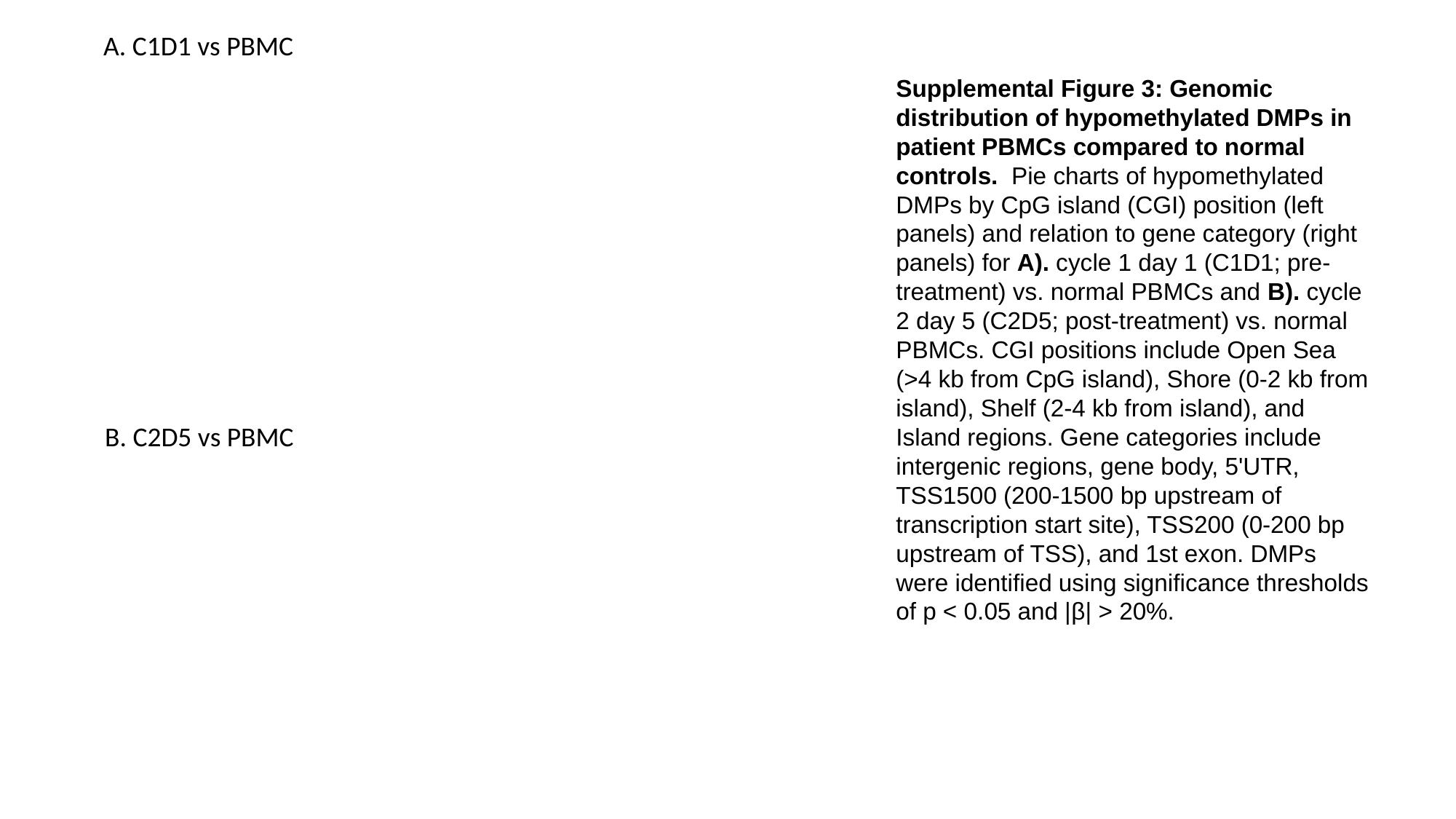

A. C1D1 vs PBMC
Supplemental Figure 3: Genomic distribution of hypomethylated DMPs in patient PBMCs compared to normal controls. Pie charts of hypomethylated DMPs by CpG island (CGI) position (left panels) and relation to gene category (right panels) for A). cycle 1 day 1 (C1D1; pre-treatment) vs. normal PBMCs and B). cycle 2 day 5 (C2D5; post-treatment) vs. normal PBMCs. CGI positions include Open Sea (>4 kb from CpG island), Shore (0-2 kb from island), Shelf (2-4 kb from island), and Island regions. Gene categories include intergenic regions, gene body, 5'UTR, TSS1500 (200-1500 bp upstream of transcription start site), TSS200 (0-200 bp upstream of TSS), and 1st exon. DMPs were identified using significance thresholds of p < 0.05 and |β| > 20%.
B. C2D5 vs PBMC
